# Fixational eye movements enable robust edge detection

**DOI:** 10.1101/2022.05.30.493986

**Authors:** Lynn Schmittwilken, Marianne Maertens

## Abstract

Human vision relies on mechanisms that respond to luminance edges in space and time. Most edge models use orientation-selective mechanisms on multiple spatial scales and operate on static inputs assuming that edge processing occurs within a single fixational instance. Recent studies, however, demonstrate functionally relevant temporal modulations of the sensory input due to fixational eye movements. Here we propose a spatiotemporal model of human edge detection which combines elements of spatial and active vision. The model augments a spatial vision model by temporal filtering and shifts the input images over time mimicking an active sampling scheme via fixational eye movements. The first model test was White’s illusion, a lightness effect that has been shown to depend on edges. The model reproduced the spatial-frequency-specific interference with the edges by superimposing narrowband noise (1-5 cpd), similar to the psychophysical interference observed in White’s effect. Second, we compare the model’s edge detection performance in natural images in the presence and absence of Gaussian white noise with human-labeled contours for the same (noise-free) images. Notably, the model detects edges robustly against noise in both test cases without relying on orientation-selective processes. Eliminating model components, we demonstrate the relevance of multiscale spatiotemporal filtering and scale-specific normalization for edge detection. The proposed model facilitates efficient edge detection in (artificial) vision systems and challenges the notion that orientation-selective mechanisms are required for edge detection.

## 1 Introduction

Edges are important features in the environment because they demarcate surface and object boundaries. An edge is a luminance discontinuity which is localized in space and/or time and extends in one direction. When edges are removed from the visual input, perception becomes less sensitive or even fades away. Replacing a luminance step by a ramp renders the luminance difference between the plateaus less visible (O’Brien, 1958). Similarly, smoothing the boundaries of geometrical shapes or stabilizing the retinal input via gaze-contingent displays leads to perceptual fading (Poletti & Rucci, 2010; Troxler, 1804). Less extreme cases of partial fading can be induced with masking (Betz et al., 2015a; Paradiso & Nakayama, 1991; Salmela & Laurinen, 2009) and contour adaptation experiments (Anstis, 2013; Betz et al., 2015b). In all these cases, spatial or temporal interference with edge-sensitive mechanisms leads to an impaired perception of surfaces and image structures.

The phenomenological importance of edges coincides with physiological mechanisms in the visual system that respond to luminance discontinuities in space and time. Neurons in the retina and thalamus with their concentric center-surround receptive fields are suited to detect edges and so are simple cells in V1 which are selective to spatial frequency and orientation (De Valois et al., 1982; Hubel & Wiesel, 1962, 1968; Parker & Hawken, 1988). This is why their function is emulated in the so-called *standard spatial vision model* (**¡**)e.g.¿schuett2017, where they are implemented as (oriented) filters at different spatial scales. Standard spatial vision models have been developed to predict human contrast sensitivity, but have also successfully been used to predict human edge sensitivity. However, despite much effort either including oriented filters at different spatial scales (Elder & Sachs, 2004; Georgeson et al., 2007) or computing additional orientation-selective features in the filtered image (such as zero-crossings in the derivatives) (Canny, 1986; Elder & Zucker, 1998; Marr & Hildreth, 1980; Watt & Morgan, 1985), it has proven difficult to robustly detect edges across different scenarios (Carandini et al., 2005).

The standard spatial vision model relies exclusively on spatial information assuming that the modeled processes occur within a single fixation and hence under static viewing. However, under natural fixation conditions, humans constantly move their eyes. These so-called fixational eye movements (FEMs) are involuntary and are composed of fast and large microsaccades and slow and small ocular drift motions (Ditchburn & Ginsborg, 1953; Ratliff & Riggs, 1950). Tapping the maximum potential of eye tracking technology, it was demonstrated empirically that ocular drift is crucial for high acuity vision (Boi et al., 2017; Intoy & Rucci, 2020; Ratnam et al., 2017) and might serve to emphasize luminance discontinuities (Rucci & Victor, 2015), i.e. edges, even in the absence of orientation-selective mechanisms (Kuang et al., 2012).

Picking up on these observations, here we combine elements from active vision and spatial vision, and propose a spatiotemporal model of human edge detection. We investigate 1) whether an active sampling of the visual input via fixational eye movements facilitates human edge detection, and 2) whether it is possible to extract edges in time and space without orientation-selective mechanisms. Our approach was as follows. We take a standard spatial vision model with unoriented Difference-of-Gaussians (DoG) filters at multiple scales and augment it with a temporal bandpass filter and subsequent temporal integration. As model input, we use a sequence of images where each time frame is sampled via ocular drift and hence a slightly shifted view of the original (static) image.

We evaluate model performance in two test cases. The first test case is the spatial-frequency-specific effect of narrowband noise on White’s illusion. Figure 1 illustrates White’s illusion in the absence and presence of narrowband noise of different center frequencies. In the original noise-free stimulus (Fig. 1A), two equiluminant gray patches appear differently bright when placed on either the dark or light bar in the carrier grating. This lightness difference between the target patches is virtually absent when the stimulus is superimposed with narrowband noise with center spatial frequencies between 1 to 5 cpd (Fig. 1C; Betz et al., 2015a; Salmela and Laurinen, 2009). As is evident from the Figure, this type of noise also effectively interferes with the vertical low contrast edges of the target (compare Fig. 1B, C and D). It has therefore been concluded that the reduction in lightness is mostly attributable to the effective masking of those edges by narrowband noise of intermediate spatial frequency (Salmela & Laurinen, 2009). Although this effect is not a direct measure of human edge perception, we think it is a useful test case for an edge detection model, because (1) it probes edge processing in a highly controlled setting, (2) it shows the relevance of different spatial frequencies for human edge detection (Elder & Sachs, 2004; Elder & Zucker, 1998), and (3) it has been shown to pose a challenge for various multiscale vision models (Betz et al., 2015a).

**Figure 1:**
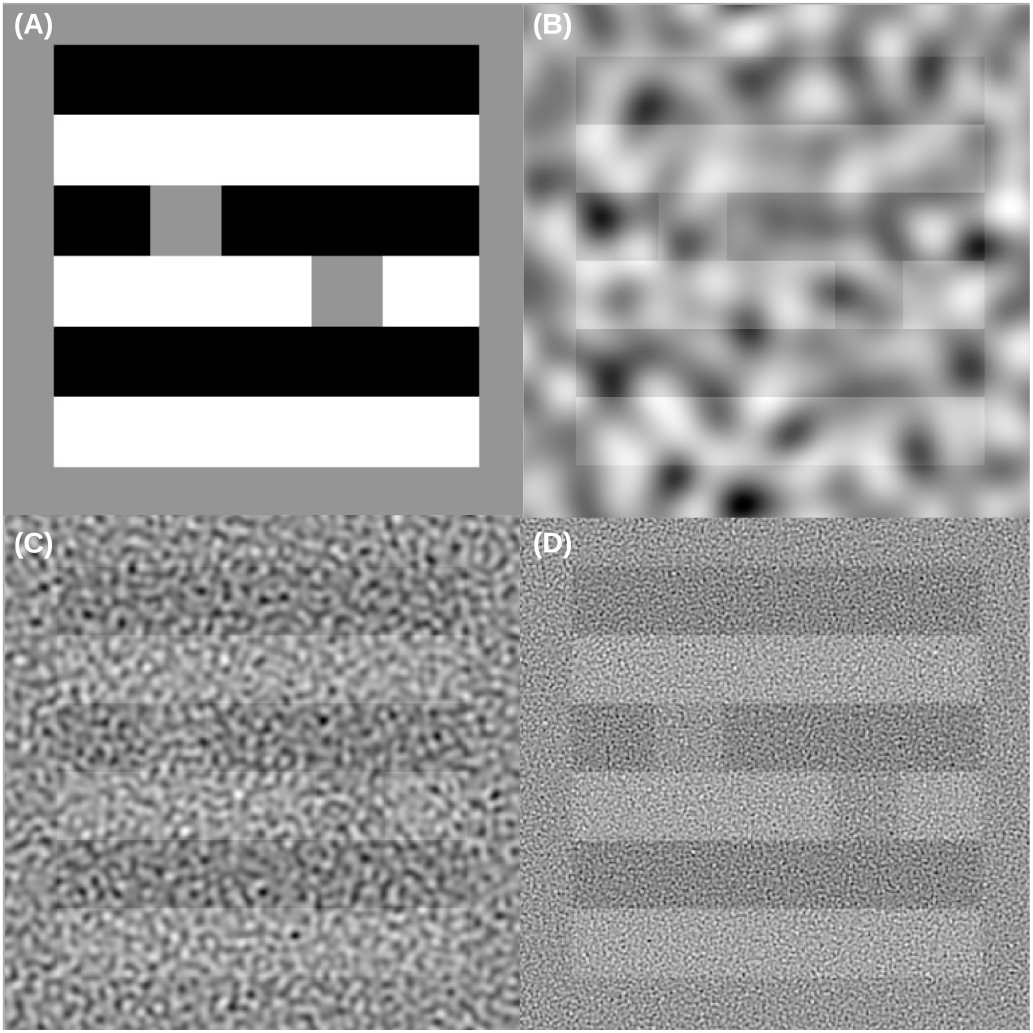
Narrowband noise between 1-5 cpd interferes with White’s effect. (A) A gray patch on a black bar is perceived as lighter than a gray patch on a white bar even though they are equiluminant (= White’s effect). (B) White’s stimulus masked with 0.58 cpd noise, (C) 3 cpd, (D) 9 cpd assuming a stimulus size of 5 cm at a viewing distance of 30 cm. White’s effect is reduced in (C). Human subjects reported that edges are hardly visible in (C). For demonstration purposes the stimulus contrast is higher in the Figure than in the actual test stimuli.

In the second test case, we compare the model’s edge detection performance to human contour detection in natural images which has become a standard comparison in computer vision. Complementary to the first test case, this investigates how the model performs under more realistic and less controlled conditions. There are several openly available benchmark datasets which contain high resolution images of different categories and so-called ground truth contour maps drawn by human observers (Fig. 2; Grigorescu et al., 2003; Martin et al., 2001). To derive these maps, observers are instructed to trace the outlines of objects and not individual edges, which is why we refer to these maps as contour rather than edge maps. To test the model’s contour detection performance, we correlated the model output with the ground truth contour maps from the Contour Image Database (Grigorescu et al., 2003). We compared how the model performed for images with and without superimposed Gaussian white noise, because edge detection in white noise is a challenge for artificial systems whereas it has relatively little effect on human perception (e.g. Burgess et al., 1981; Goris et al., 2008; and compare Fig. 2 A and B).

**Figure 2:**
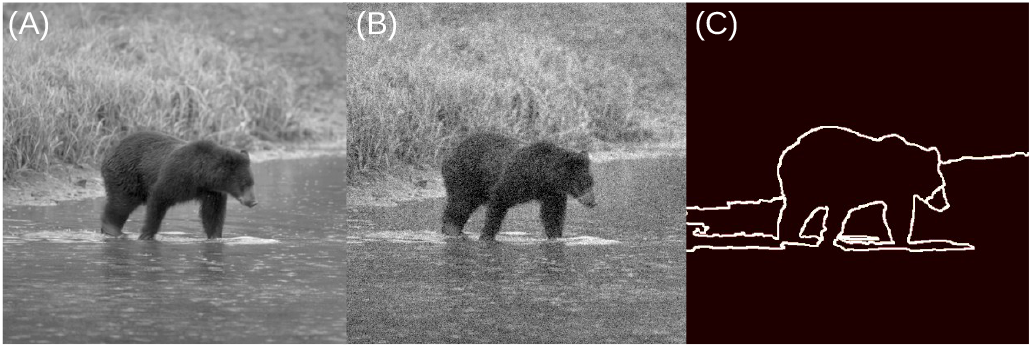
(A) Example natural image taken from the Contour Image Database (Grigorescu et al., 2003). (B) We added Gaussian white noise to the images to test the robustness of the model against noise. (C) Human-drawn contour map provided with the Contour Image Database (Grigorescu et al., 2003) which represents human contour detection performance and which we use to evaluate model performance in both noise conditions.

To anticipate, the model reproduced the spatial-frequency-specific interference of narrowband noise on the edge representations underlying White’s illusion and robustly detected edges in natural images. Edge signals emerged in the spatiotemporal filter responses of the model as a consequence of actively sampling the visual input. Orientation-selective mechanisms or other directional features were not required.

## 2 Methods

### 2.1 Active spatiotemporal edge detection model

Figure 3 shows a schematic overview of the model ^1^. To mimic the active nature of visual processing during fixations, we sample images from slightly different viewpoints as they would occur as a consequence of ocular drift. The resulting dynamic image sequence is illustrated in Figure 3B. We simulate drift as Brownian motion (Kuang et al., 2012) with a temporal frequency of *f* = 100*Hz*. This samples the temporal range that the visual system is most sensitive to (Derrington & Lennie, 1984; Zheng et al., 2007), while keeping the computational cost low. We define the time period *T* = 0.2*s* as this was proposed as minimal fixation duration (Salthouse & Ellis, 1980). We set the diffusion coefficient 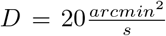 to fit the magnitude of recorded drift motions (Kuang et al., 2012). The resulting input videos consists of *T · f* + 1 = 21 frames.

**Figure 3:**
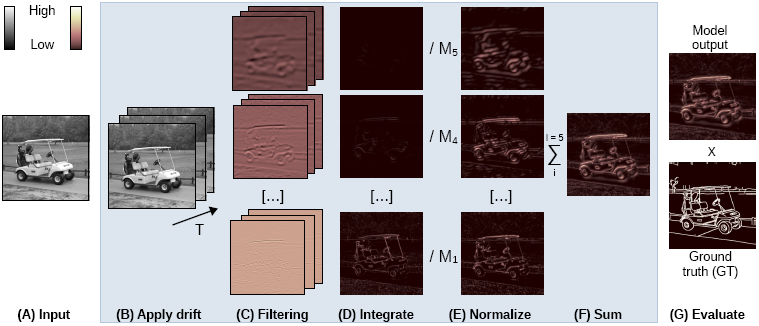
Model structure. We simulate the active sampling of the visual system within one fixation by applying drift (B) to the input (A). We then filter the dynamic input with multiple spatial and one temporal filter (C). Each row illustrates the filtered responses at different spatial scales *i*. We extract edges via temporal integration at each spatial scale (D). After normalization by the global means *M*_*i*_ (E), we sum the signals into the final model output (F). We quantify model performance as correlation between the aligned model output and the ground truth (G).

This dynamic input is then fed to the spatiotemporal vision model which is built upon the standard components of early vision models: linear filtering at multiple scales followed by non-linear response normalization within and integration across spatial scales (Carandini et al., 1997; Heeger, 1992; Schütt & Wichmann, 2017). We filter the dynamic input in space and time (Fig. 3C). Inspired by spatial tuning properties of simple cells (De Valois et al., 1982), we use five spatial DoG filters *G*_*i*_(*f*_*x*_, *f*_*y*_) with peak spatial frequencies (SFs) between 0.62 and 9.56 cpd in octave intervals (Fig. S1A-E) defined in the frequency domain as

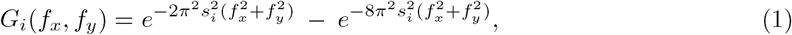

where *f*_*x*_ and *f*_*y*_ denote the SFs in cpd and *s*_1*-*5_ = [0.016, 0.032, 0.064, 0.128, 0.256] deg controls the spatial scale of the DoG filters.

The temporal bandpass filter *H*(*ω*) was fitted to the temporal tuning properties of macaque simple cells reported in (Zheng et al., 2007) with a peak frequency of 9.52 Hz and no sensitivity to static inputs (Fig. S1G). It is defined in the frequency domain as

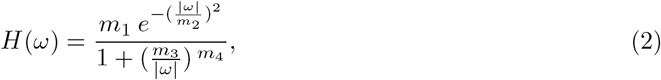

where *ω* denotes the temporal frequencies in Hz, *m*_2_ = 22.9, *m*_3_ = 8.1, *m*_4_ = 0.8, and *H*(*ω* = 0) = 0. Since *m*_1_ was negligible here, we set *m*_1_ = 1 (instead of *m*_1_ = 69.3).

To spatiotemporally filter the input video *S*(*x, y, t*), we perform

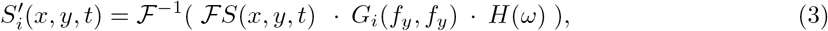

where 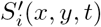 denotes the filtered video at each spatial scale *i*, ℱ denotes the Fourier transform and ℱ^−1^ denotes the inverse Fourier transform.

To temporally integrate the filtered videos 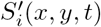, we compute the squared mean separately at each scale *i* (Fig. 3D). Then, we normalize the integrated signals 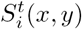 by their mean activation *M*_*i*_ (Fig. 3E). In the final step (Fig. 3F), we sum the normalized signals over all scales into the model output according to

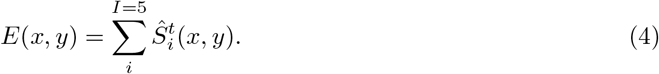

To avoid boundary effects, we crop the outer 0.5 deg of the edge map *E*(*x, y*). To quantify edge detection performance (Fig. 3G), we correlate the model output *E*(*x, y*) with a ground truth template (Sebastian et al., 2017). We define the ground truth template as matrix of the same size as the model output with non-zero entries at the location of the edges and zero entries everywhere else. Since the edge locations in *E*(*x, y*) slightly vary depending on the exact drift trace, we align *E*(*x, y*) and the ground truth edge template to maximize cross-correlation before we calculate the pixel-wise Pearson correlation.

### 2.2 Eliminating model components

To test which model components are crucial for robust edge detection, we systematically eliminated model components (Fig. 3C-E). We label the full model as ST-M-N (SpatioTemporal filtering - Mean - Normalization) and label the control models accordingly.

- **S-M-N (Spatial filtering - Mean - Normalization)**: This model omits temporal filtering. It performs multiscale spatial filtering (S), temporal integration via the squared mean (M) and normalization within scales (N).
- **ST-M (Spatiotemporal filtering - Mean)**: This model omits the normalization. It is reduced to spatiotemporal filtering (ST) followed by temporal integration via the squared mean (M).
- **T-M (Temporal filtering - Mean)**: This model omits multiscale spatial filtering. It is reduced to temporal filtering (T) followed by temporal integration via the squared mean (M). Without multiple scales, scale-specific normalization is not required.
- **Canny**: We also compare the model with an optimized Canny edge detector (Canny, 1986) because of its wide use in computer vision. Optimization was required for each test case individually to achieve good performance. Technical implementation details are reported in the appendix.

#### Computing the variance of the dynamic input

Mathematically, computing the variance of a signal over time is analogous to filtering the signal with a temporal filter that is insensitive to static inputs, and temporally integrating via the squared mean. To show this equivalence, we introduce a second group of control models in which we substitute the temporal filtering (Fig. 3C) by a variance operation in the integration step (Fig. 3D). Analogous to the first group of control models, we progressively eliminate model components to identify the ones crucial for robust edge detection.

- **S-V-N (Spatial filtering - Variance - Normalization)**: Counterpart to full model (ST-M-N). It substitutes temporal filtering (S) and integration via the squared mean by a variance operation (V). It involves a non-linear normalization at each spatial scale (N).
- **S-V (Spatial filtering - Variance)**: Counterpart to ST-M model. It replaces temporal filtering (S) by a variance operation (V) and omits normalization.
- **V (Variance)**: Counterpart to T-M model. It perform no filtering, but directly computes the variance (V) of the dynamic input over time.

### 2.3 Test case 1: The spatial-frequency-specific effect of narrowband noise

First, we tested whether the model can predict the SF-specific interference of narrowband noise with the edge representations in White’s stimulus. When White’s stimulus (Fig. 1A) is superimposed with narrowband noise of about 1-5 cpd, the targets are hardly visible and the lightness effect is substantially reduced (see Fig. 1B-D). In psychophysical experiments, observers’ perceived lightness of each target patch was measured in a matching task (Betz et al., 2015a; Salmela & Laurinen, 2009). The effect of noise on perceived lightness was quantified as the difference in matched lightness for the target placed on the light vs the dark bar. Target lightness was measured for different grating frequencies and the strength of the illusion was selectively reduced for noise center frequencies of 1-5 cpd (Fig. 4B).

**Figure 4:**
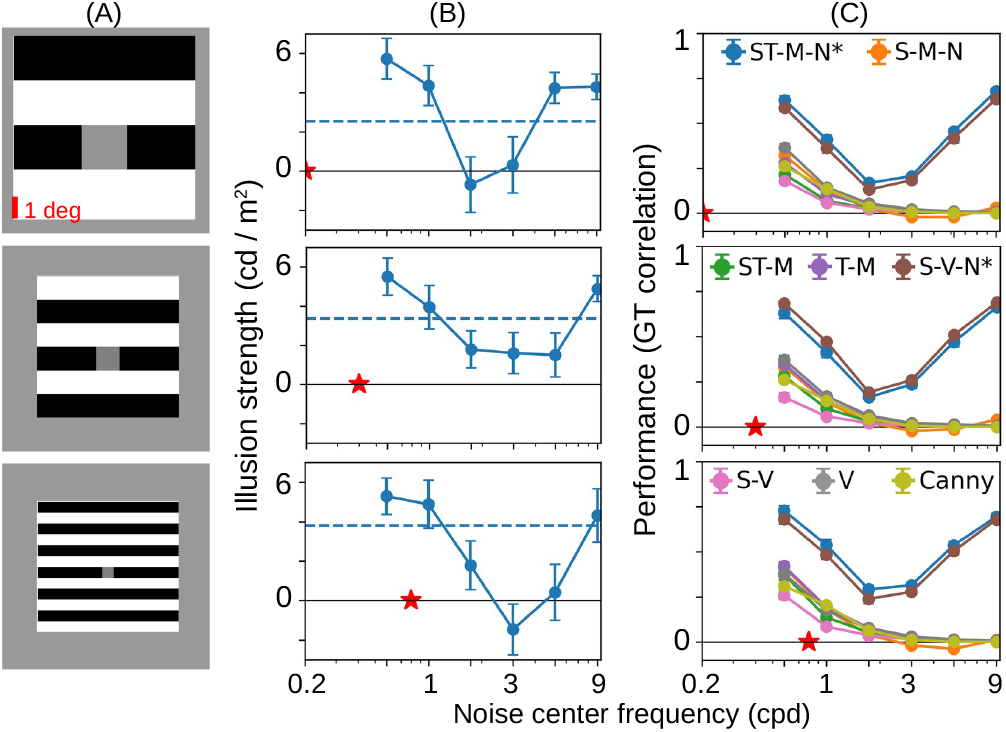
Human and model performance in test case 1. Different rows show White’s stimulus with different grating frequencies (A) and corresponding human (B) and model (C) results. (B) Psychophysically measured illusion strength (n=11) as function of noise center frequency (x-axis). Error bars indicate standard errors of the mean. Red stars indicate the spatial frequency of the carrier grating. Dashed lines indicate illusion strength for stimuli without noise. (C) Effect of narrowband noise on the edge detection performance of all models (n=10 trials).

Here we test whether the model’s edge detection performance is also specifically impaired by narrowband noise with center frequencies between 1-5 cpd in comparison to noise with center frequencies outside this range. We compare the model’s edge detection performance to the psychophysical illusion strength (Betz et al., 2015a) assuming that the reduction in illusion strength is predominantly caused by interference with the edges. The question how exactly perceived lightness, and thus White’s effect, is derived from the representation of the edges is an open research question and beyond the scope of the proposed model. It most likely involves additional mechanisms such as edge integration, iso-orientation suppression, figure-ground segmentation, or filling-in (**¡**)e.g.¿betz2015b, domijan2015, grossberg1988.

We re-implemented the stimuli used in Betz et al. (2015a) that depict single-target White’s stimuli (Fig. 4A) and superimposed them with narrowband noise. Stimuli were comprised of horizontal square wave gratings (Michelson contrast = 0.05) with black and white bars and a single gray patch. We used three versions of White stimulus with different grating frequencies (Fig. 4A). The gratings with low, medium and high SFs contained 4, 6 and 12 bars, corresponding to SFs of 0.2, 0.4 and 0.8 cpd. Total stimulus and patch sizes were different for different SFs. The low SF grating was 10.2×10.2 deg with patch size 2.55×2.55 deg, the medium SF grating was 7.65×7.65 deg, with patch size 1.28×1.28 deg, and the high SF grating was 7.50×7.50 deg, with patch size 0.63×0.63 deg. The stimuli were masked with bandpass-filtered white noise with one octave SF bandwidth. Corresponding to the psychophysical design, we used six noise masks with random instantiations and noise center frequencies between 0.58 to 9 cpd in logarithmic steps. All noise masks had a size of 16×16 deg with RMS contrasts of 0.2 (standard deviation divided by mean). Since ocular drift evokes small shifts of the input, we required a high stimulus resolution of 40 pixels per degree. Afterwards, we fed the stimuli to the model and computed the model response for each stimulus. We created ground truth templates with the edge locations and magnitudes of the noise-free stimuli. Edge thickness was 0.1 deg as this roughly matches the edge thickness generated by the models. Model performance was quantified as Pearson correlation between the ground truth edge templates and the aligned model outputs.

### 2.4 Test case 2: Contour detection in natural images

To test the model under less artificial conditions, we evaluated its contour detection capacities on natural images. Images were taken from the Contour Image Database (Grigorescu et al., 2003) which contains 40 grayscale images (512×512 pixels) of textured scenes depicting man-made objects or animals in their natural habitat. Each image is accompanied by a human-drawn contour map which we use as ground truth to evaluate the different model performances (see Fig. 2C). The original purpose of the database was to evaluate the contour detection performance of different algorithms while suppressing texture edges. Observers were thus instructed to trace all contours but not necessarily all edges (Grigorescu et al., 2003). The ground truth maps therefore reflect human contour rather than human edge perception.

We preprocessed the images before feeding them into the models. We converted the image sizes from pixels to degrees visual angle with a resolution of 40 pixels per degree. This resulted in image sizes of 12.8×12.8 degree. Additionally, we normalized the pixel intensities between 0 and 1. To quantify model performance relative to human performance, we correlated the model outputs with the human-drawn contour maps from the database. We again increased edge thickness in the ground truth maps to 0.1 deg to roughly match the edge thickness generated by the models.

To evaluate the models’ robustness against noise, we tested each of the different models on the images from the database in the presence and absence of Gaussian white noise. To keep the overall brightness between images with and without noise comparable (average difference in mean brightness = 0.0014), we used noise with *µ*_*noise*_ = 0 and *σ*_*noise*_ = 0.1 and cropped pixel intensities smaller than 0 and larger than 1 (values were substituted by 0 and 1, respectively). We used Gaussian white noise because it challenges the edge detection capacities of artificial systems, while it has been shown to be relatively ineffective for human perception (Burgess et al., 1981; Goris et al., 2008). This can be informally confirmed by comparing Figures 2A and B. Model performance was quantified as Pearson correlation between the human-drawn contour maps and the aligned model outputs. Human contour maps were only available for images without noise (Grigorescu et al., 2003).

## 3 Results

### Test case 1: The spatial-frequency-specific effect of narrowband noise

Figure 4B shows the psychophysical data for White’s effect as a function of the center frequency of the narrowband noise (data from Betz et al., 2015a). Analogously, Figure 4C shows the edge detection performance of the models. The full model (ST-M-N, blue curve) and its variance counterpart (S-V-N, brown curve) show the highest and very similar performance. More importantly, they the SF-specific effect of narrowband noise on White’s effect as the edge representations of these two models are most impaired for noise center frequencies between 1-5 cpd. All other models show an overall lower performance do not exhibit the SF-specific dip. Their performance rather decreases with increasing noise center frequency. This is also the case for the Canny edge detector (lime curve) despite being optimized to reproduce the psychophysical effect.

### Test case 2: Contour detection in natural images

Figure 5 plots the contour detection performance of the models quantified as Pearson correlation between model outputs and human contour maps from the database (Grigorescu et al., 2003). The full ST-M-N model (blue curve), its variance counterpart S-V-N (brown curve) and the optimized Canny edge detector (lime curve) perform better than all other models in the absence and in the presence of Gaussian white noise (compare Fig. 5A and B). The models without multiscale spatial filtering (T-M, V; lilac and gray curves) or without normalization (ST-M, S-V; green and pink curves) were most affected by the noise. The performance of the full ST-M-N model, the ST-M and T-M model are close to the performances of their variance counterparts S-V-N, S-V and V.

**Figure 5:**
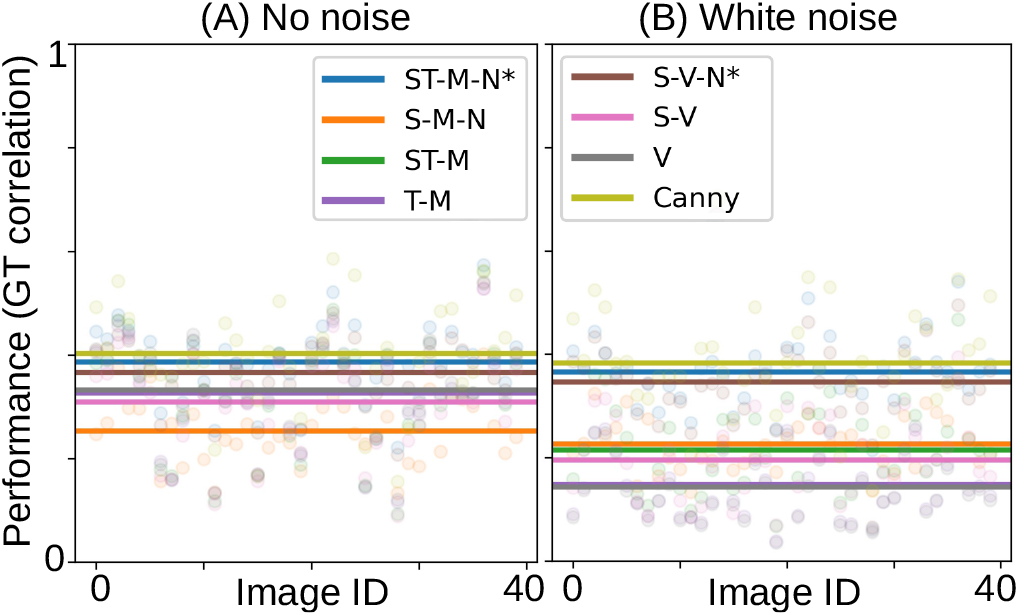
Models’ contour detection performance in natural images in the (A) absence and (B) presence of Gaussian white noise. Lines indicate average model performance over all 40 images from the database (N=10 trials). The full ST-M-N model and its variance counterpart S-V-N are marked with an asterisk. In panel A, the lines indicating performance of the ST-M (green), T-M (lilac) and V (gray) models, lie on top of each other. In panel B, the lines indicating performance of the T-M (lilac) and V (gray) models, lie on top of each other. For results, see text.

Figure 6 shows the model outputs (or model activations) before correlation with the human-drawn contour maps. In the absence of noise, almost all models respond to object outlines (e.g. bear), but also respond to textural structures such as grass or reflections on the water. The purely spatial S-M-N model generally produces low activation for contours, but high activation for features such as the legs of the bear. The S-M-N model demonstrates that additional, direction-selective features are required to extract edges in purely spatial models.

**Figure 6:**
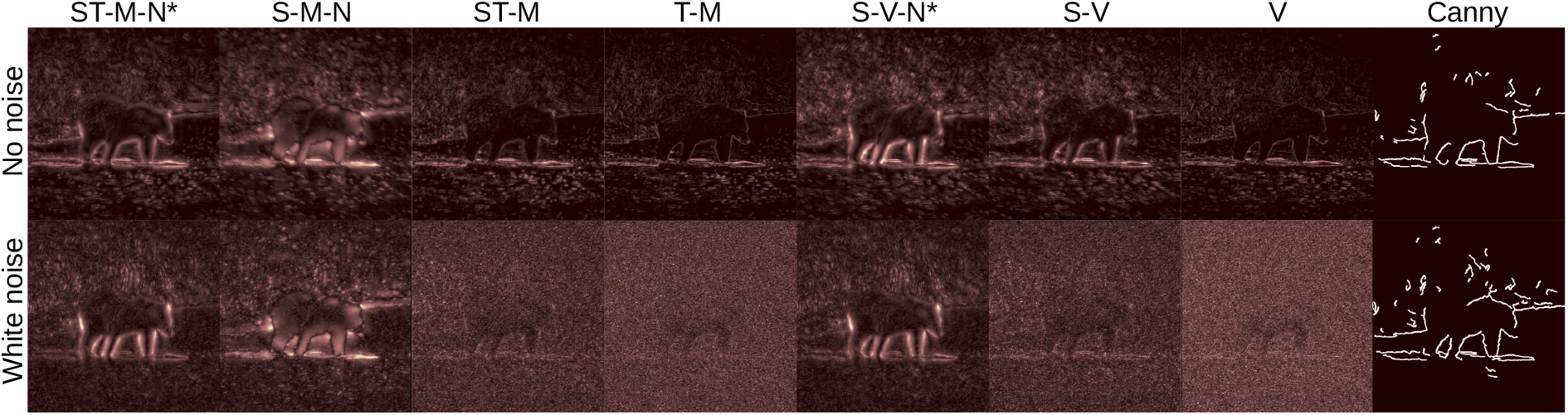
Model outputs for example input image from Contour Image Database (Grigorescu et al., 2003) shown in Figure 2 for the no-noise condition (top row) and the Gaussian white noise condition (bottom row). For results, see text.

Model performances vary substantially between different input images (compare individual data points in Fig. 5). Visual inspection of the model outputs revealed that models perform poorly on images that contain many high-contrast textures (see examples in Fig. 6). This is not too surprising, since we compare the output of edge detection models to human-drawn contour maps. ”Falsely” detected edges were to be expected as not all edges form part of a contour.

## 4 Discussion

The aim of this study was to investigate whether a spatiotemporal early vision model could successfully extract edges in short fixational instances when the input is actively sampled. In particular, we tested whether a spatiotemporal strategy enables edge detection in the absence of orientation-selective mechanisms.

Standard spatial vision models come in different flavors, but their basic components are SF- and orientation-selective units followed by non-linear response normalization and integration across channels (Carandini et al., 1997; Geisler & Albrecht, 1995; Heeger, 1992). They can account for many perceptual phenomena such as contrast sensitivity, edge detection, and brightness perception (e.g. Blakeslee and McCourt, 1999; Georgeson et al., 2007; Schütt and Wichmann, 2017). Spatial vision models operate on static inputs assuming that the eyes are mostly still during fixations, and that FEMs, if anything, introduce noise to the sensory input. Over the last two decades, however, evidence has accumulated that FEMs play an active role in visual processing and might emphasize edge signals in the visual input (Kuang et al., 2012; Rucci & Victor, 2015).

Inspired by these findings, we tested whether ocular drift plays a role in human edge processing by sampling the visual input in a characteristic way during fixations. We tested the behavior of a standard spatial vision model with unoriented filters that we (1) extended by a temporal dimension, and (2) fed with an actively-sampled sensory input. Such a model is reminiscent of existing spatiotemporal models of human motion perception (e.g. Adelson and Bergen, 1985) but has not been applied to static images at the time scale of fixations. Our simulations show that edge signals emerge in the spatiotemporal filter responses of the proposed model as a consequence of the active sampling. The full model (ST-M-N) outperformed an optimized Canny edge detector as well as the purely spatial (S-M-N) and the purely temporal (T-M) control models. It reproduced the spatial-frequency-specific interference of narrowband noise of 1-5 cpd with the edge representations underlying White’s illusion, and it robustly detected edges in natural images in the presence and absence of Gaussian white noise.

In addition, we show that temporal filtering and integration can be substituted by computing the variance of the time-varying filter responses. Computing the variance is more efficient, because it requires a lower temporal sampling of the visual input than the temporal filtering operation. It has been suggested before that the temporal response variations of cells at the front-end of the visual system can be used for edge detection. Hongler et al. (2003) described the purely temporal *resonant retina* model which exploits small camera vibrations to detect edges (see also Prokopowicz and Cooper, 1995). A related idea is exploited in neuromorphic event-based vision systems which only register changes over time and thus efficiently code input features such as edges (Gallego et al., 2020). The resonant retina and the event-vision ideas are most similar to our control model V, which was however prone to high spatial frequency noise in both test cases, and performed worse than the model proposed here. We will discuss the relevance of multiscale spatial filtering and response normalization below.

### Edge detection without orientation-sensitive mechanisms

Many neurons in V1 respond preferentially to oriented bars of various width and length, which is why V1 has been ascribed the role of edge analysis of the visual input (Hubel and Wiesel, 1962, 1968; but see Carandini et al., 2005). In particular simple cells have served as inspiration for models of human edge processing. Analogously to simple cells, these models use linear filters at multiple spatial scales that are sensitive to orientation. There are two sub-categories of simple-cell-inspired edge models: (1) models that use oriented filters (e.g. Gaussian derivative filters) to detect edges (Georgeson, 1992; Georgeson et al., 2007), and (2) models that use unoriented filters (e.g. DoG or Gaussian filter) and detect edges by computing additional directional features (e.g. zero-crossings or directional derivatives) of the filtered outputs (Canny, 1986; Elder & Zucker, 1998; Marr & Hildreth, 1980; Watt & Morgan, 1985). What all these models have in common is some dedicated orientation-sensitive mechanism (oriented filters, zero-crossing or directional derivatives) which is used to extract edge signals from the visual input ^2^.

The present model extracts edge signals without dedicated orientation-selective processing because the differential signal at luminance discontinuities is conveyed in time. Temporal filtering of an actively sampled input is sufficient to convert the temporal variations of the input signal into discontinuities in space (Fig. 7). This may come as a surprise at first, since particularly their orientation-selectivity made simple cells popular as edge detectors. However, it has previously been shown that model neurons with circularly symmetric receptive fields, and hence no orientation preference, emphasize edges in an image (Kuang et al., 2012). The present model is still inspired by properties of neurons in V1. It performs multiscale spatial filtering of the visual input at each location (Campbell & Robson, 1968) and it emulates spatial (De Valois et al., 1982) and temporal (Zheng et al., 2007) sensitivities of simple cells. Also, there is physiological data showing that a substantial number of simple cells are not tuned to orientation (Ringach et al., 2002; Talebi & Baker, 2016). These cells have shorter response durations than orientation-selective cells, are highly reliable and show no direction-selectivity (Talebi & Baker, 2016). Such non-oriented simple cells seem to be well suited to extract edge signals from the visual input when considering drift motions.

**Figure 7:**
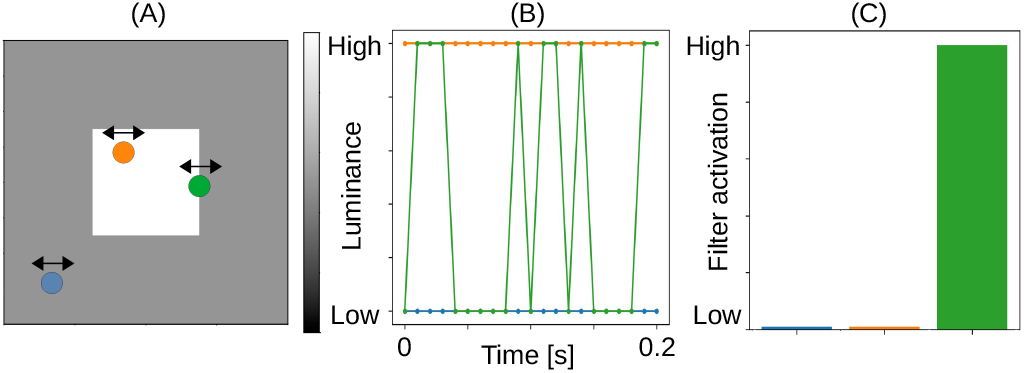
Consequence of ocular drift illustrated for an example input image and for units at three different locations in (A). The arrows indicate location shifts that results from ocular drift. (B) The resulting dynamically-sampled luminance signal is flat for units at homogeneous image regions (orange and blue) but fluctuates over time for units close to luminance edges (green). (C) A temporal filter that only responds to fluctuating input signals, will not be activated by flat luminance signals (orange and blue) but will generate large activations for the fluctuating luminance signal (green).

Unoriented filters are computationally more efficient for edge detection (Marr & Hildreth, 1980). Detecting edges via ocular drift might reduce the number of filters tuned to both spatial frequency and orientation. In order to achieve orientation sensitivity in further processing stages, the output of the proposed model could still be processed by orientation-selective cells but these could be tuned to a much narrower spatial scale.

### Relevance of multiscale filtering and non-linear normalization

There is general agreement that the visual system uses multiscale processing to extract edges (Elder & Zucker, 1998; Watt & Morgan, 1985). The reasoning for this is two-fold. Edges in the visual input come in different sizes, at different distances, and at different sharpness and hence differ in their amount of blur. Filters at different spatial scales seem to be a proper response to this variety. A multiscale approach also proves beneficial when edges have to be detected in the presence of noise. Our results corroborate the importance of multiscale filtering for edge detection, because performance substantially dropped in the presence of noise for the purely temporal models (T-M, V model) and for the Canny edge detector. This is particularly evident in test case 1 (Fig. 4C), presumably because the contrast of the noise was comparatively large relative to that of the grating.

A multiscale filtering system differs from a single-scale system, only if there is subsequent scale-specific processing. Our simulations suggest that it is the non-linear normalization within each spatial scale which is crucial for robust edge detection. This became evident from the model responses to the noisy images in test case 2, where performance substantially dropped for control models without normalization (Fig. 5; ST-M, T-M, S-V, V). We observed that without scale-specific normalization, high spatial frequency contents in the images dominated the model outputs (Fig. 6; ST-M, T-M, S-V, V).

Here we implemented a divisive normalization scheme which was suggested to account for response non-linearities of simple cells in macaque V1 (Carandini et al., 1997; Geisler & Albrecht, 1995; Heeger, 1992). This follows the tradition of linear multiscale filtering followed by a divisive response normalization which is routinely used in early vision models (Malik & Perona, 1990; Schütt & Wichmann, 2017; Teo & Heeger, 1994; Watson & Solomon, 1997). Nonetheless, it should be mentioned that the global normalization scheme employed in the model is neurally not plausible, because it would require lateral or feedback connections across the whole visual field (c.f. discussion in Robinson et al., 2007). A more realistic normalization scheme should be local and it should consider that cells with similar spatial frequency preferences interact more (Issa et al., 2000). The main reason for implementing a global normalization scheme was that it was required to reproduce the psychophysical results in test case 1 and that it was simple. Edge extraction in (noisy) natural images, as in test case 2, was not sufficient to further constrain the normalization scheme of the model. We need more psychophysical data from experiments that specifically probe human edge sensitivity in the presence of various types of noise in order to constrain a more biologically-plausible normalization scheme.

### 4.1 Specificity of temporal filtering

The model’s edge detection capacities can be attributed to the temporal filter which responds only to dynamically changing parts in the input. To illustrate what this means we have visualized the consequence of ocular drift for homogeneous image regions and for a luminance edge in Figure 7. Close to the edge, ocular drift causes small image shifts between dark and bright areas and hence causes the dynamically-sampled signal to fluctuate over time (green location in Fig. 7). A temporal filter tuned to non-zero temporal frequencies will respond to these fluctuations. The dynamically-sampled signal in homogeneous image regions will stay flat over time (orange and blue locations in Fig. 7). The temporal filter will not respond to these static inputs. This model feature is in agreement with the finding that the visual system is most responsive to luminance transients (Derrington & Lennie, 1984; Hawken et al., 1996). Our results show that the specific properties of the temporal filter are not crucial to detect edges. We can even substitute the temporal filtering and subsequent integration by the variance operation. The main difference between the two approaches is that the temporal filter weighs the contributions of individual spatial frequency components differently whereas the variance does not.

It has been shown that the temporal transients that result from ocular drift specifically enhance the high spatial frequency contents in the input (Rucci et al., 2018; Rucci & Victor, 2015). In the proposed model, we neither observe an additional effect of temporal filtering (compared to the variance), nor an enhancement of high spatial frequencies. This is because we apply a spatial-scale-dependent normalization scheme which equalizes the contribution of each spatial scale in the final model output. Control models without spatial-scale-dependent normalization produced model outputs dominated by high spatial frequency contents (compare Fig. 5; ST-M, T-M, S-V, V). This is to be expected because ocular drift enhances high spatial frequencies. Since our simulations suggest that robust edge detection requires both e multiscale filtering as well as a scale-dependent normalization, it would be interesting to see whether and how a more biologically-plausible normalization scheme could solve this apparent discrepancy.

### 4.2 Limitations of test cases

To our knowledge, there is no standardized approach to test models of human edge perception. Possible test cases have been blur coding (Elder & Zucker, 1998; Georgeson et al., 2007), Mach bands (Watt & Morgan, 1985), edge localization (McIlhagga, 2018) and edge polarity discrimination in noise (Elder & Sachs, 2004).

Here, we have used two test cases to evaluate model performance - one controlled and one more natural task. The first test case was the spatial-frequency-specific interference of superimposed narrowband (1-5 cpd) noise on the edge representations underlying White’s illusion. We chose this test case, because (1) it probes edge processing in a controlled psychophysical setting, (2) it shows the relevance of different spatial frequencies for human edge detection (Elder & Sachs, 2004; Elder & Zucker, 1998), and (3) it challenges multiscale vision models (Betz et al., 2015a). Although the lightness effect in White’s stimulus is an indirect measure of human edge mechanisms, there are several findings which support the assumption that the spatial-frequency-specific interference with perceived lightness can be ascribed to edge-sensitive mechanisms. As can be seen in Figure 1, and as reported by participants in Betz et al. (2015a), the edges in White’s stimulus are barely visible when superimposed with narrowband noise around 3 cpd. Contour adaptation, which explicitly interferes with edge-sensitive mechanisms, has been shown to have a strong effect on lightness phenomena (Anstis, 2013). In particular, contour adaptation of the edges orthogonal to the grating has been shown to abolish White’s effect (Betz et al., 2015b). Thus edges are critical for perceived lightness at least in White’s effect, and we also assume that the spatial-frequency-specific interference of narrowband noise with White’s illusion is a valid measure of human edge-sensitive mechanisms. To fully account for the lightness effect, i.e. its magnitude, additional mechanisms such as edge integration, iso-orientation suppression, figure-ground segmentation, or filling-in (**¡**)e.g.¿betz2015b, domijan2015, grossberg1988 need to be considered.

The second test case was contour detection in natural images in the presence and absence of Gaussian white noise. We have chosen this test case to investigate to which extent the edge detection capacities of the model generalize to the more naturalistic task of human contour detection. We compared the performance of our model with the human-drawn contour maps provided in the Contour Image Database (Grigorescu et al., 2003) which can be considered an approximation of human ground truth. Human participants were specifically asked to focus on the outlines of objects and ignore more textural features for this task. In fact, it is not clear what a full human-drawn edge map (instead of contour map) would look like in natural images and how this could be tested.

Natural images differ in their spectral contents from the psychophysical stimuli used in test case 1 showing a characteristic reduction in spatial frequency components that is inversely proportional to the frequency itself. Contour detection in natural images in the absence of additional noise does not constrain human edge models well, because the models can simply extract edges as high spatial frequency contents in the input images. However, as should be clear from test case 1, the human visual system extracts edges through a rather narrow, intermediate spatial scale around 3 cpd (Fig. 4; Shapley and Tolhurst, 1973). To test the robustness of the tested models against small amounts of noise, we have therefore added the Gaussian white noise condition. We did not explicitly test it, but the small amount of Gaussian white noise did not seem to affect human contour detection much as the noise is barely visible to the human visual system (compare Fig. 2; Burgess et al., 1981; Goris et al., 2008).

## Conclusion

Existing models of early visual processes tend to focus on spatial mechanisms to account for human vision, often assuming short fixation periods in which the retinal image is stable over time. This assumption disregards the fact that our eyes are constantly moving, and that our visual system responds best to inputs that change in space and time. Here, we have proposed a spatiotemporal model of human edge processing that combines standard components of spatial vision models with an active-sampling strategy and temporal filtering. The model successfully reproduced the spatial-frequency-specific interference of narrowband noise on the edge representations underlying White’s effect and robustly detected edges in natural images with and without Gaussian white noise. In the absence of dedicated orientation-sensitive mechanisms, the model challenges the notion that orientation-selectivity is required for edge detection. Our findings corroborate the notion that human vision is an active process, and that the visual system encodes visual information efficiently in space and time.

## 5 Acknowledgments

This work was funded by the Deutsche Forschungsgemeinschaft (DFG, German Research Foundation) under Germany’s Excellence Strategy -EXC 2002/1 ”Science of Intelligence”-project number 390523135 and individual grants MA 5127/4-1 and 5-1 to M. Maertens. No commercial relationships.

## Footnotes

1 We provide a Jupyter-Notebook with the model implementation.

2 We provide a Jupyter-Notebook which implements the mechanisms employed by classical edge models.

